# Comparison of Explainable AI Models for MRI-based Alzheimer’s Disease Classification

**DOI:** 10.1101/2024.09.17.613560

**Authors:** Tamoghna Chattopadhyay, Neha Ann Joshy, Chirag Jagad, Emma J. Gleave, Sophia I. Thomopoulos, Yixue Feng, Julio E. Villalón-Reina, Emily Laltoo, Himanshu Joshi, Ganesan Venkatasubramanian, John P. John, Greg Ver Steeg, Jose Luis Ambite, Paul M. Thompson

**Affiliations:** Imaging Genetics Center, Mark and Mary Stevens Neuroimaging and Informatics Institute, Keck School of Medicine, University of Southern California, Marina del Rey, CA, United States; Multimodal Brain Image Analysis Laboratory; Translational Psychiatry Laboratory, National Institute of Mental Health and Neuro Sciences (NIMHANS), Bengaluru, Karnataka, India; University of California, Riverside, CA, United States; Information Sciences Institute, University of Southern California, Marina del Rey, CA, United States

**Author notes:** Email: {, }.

**Keywords:** Magnetic Resonance Imaging, Alzheimer’s Disease, Deep Learning, Occlusion Sensitivity Analysis, Grad-CAM

## Abstract

Deep learning models based on convolutional neural networks (CNNs) have been used to classify Alzheimer’s disease or infer dementia severity from 3D T1-weighted brain MRI scans. Here, we examine the value of adding occlusion sensitivity analysis (OSA) and gradient-weighted class activation mapping (Grad-CAM) to these models to make the results more interpretable. Much research in this area focuses on specific datasets such as the Alzheimer’s Disease Neuroimaging Initiative (ADNI) or National Alzheimer’s Coordinating Center (NACC), which assess people of North American, predominantly European ancestry, so we examine how well models trained on these data generalize to a new population dataset from India (NIMHANS cohort). We also evaluate the benefit of using a combined dataset to train the CNN models. Our experiments show feature localization consistent with knowledge of AD from other methods. OSA and Grad-CAM resolve features at different scales to help interpret diagnostic inferences made by CNNs.

## I. Introduction

Explainable AI (xAI) is crucial in medical imaging and healthcare to foster trust, transparency, and efficacy in clinical settings. With the advent of artificial intelligence in healthcare, particularly in medical imaging, AI models have shown great promise in diagnosing and subtyping neurological conditions. However, the opacity of these models poses significant challenges for their adoption in clinical practice. Clinicians need to understand and trust AI-driven decisions to integrate them effectively into patient care. xAI bridges this gap by providing insights into how AI models arrive at their conclusions, enhancing their interpretability and trustworthiness. Interpretability is not just a technical requirement but a fundamental necessity to meet the ethical and legal standards in healthcare, ensuring that decisions made by AI can be validated and audited by human experts [1].

Several techniques have been developed to explain the inner workings of AI models in neuroimaging. Grad-CAM (gradient-weighted Class Activation Mapping) [2] uses gradients of the last, or several, convolutional layers to localize critical key features for making a decision. This is particularly useful in neuroimaging [3,4] to identify specific brain regions contributing to diagnostic classification. Similarly, *saliency maps* [5,6] highlight pixels in an image that most influence the model’s prediction, improving interpretability. Whereas the former highlights regions specific to a particular class prediction, the latter gives a more general view of influential pixels regardless of the specific class. Occlusion sensitivity analysis (OSA) [7,8], by contrast, systematically occludes parts of the input image to observe changes in the model’s output, pinpointing crucial regions for decision-making. These techniques can each improve the transparency of AI systems, enhancing their value for both research and clinical settings.

Alzheimer’s disease (AD) is the leading age-related neurodegenerative disorder, accounting for approximately 70% of dementia cases worldwide [9]. Consequently, there is an urgent need to identify factors that contribute to or protect against the onset or progression of dementia. Schlemper et al. [10] proposed a novel attention gate (AG) model that learns to focus on target structures of varying shapes and sizes in medical images, improving performance on image classification and segmentation tasks while providing insights into the model’s decision-making process. A recent study [11] trained a deep CNN, based on the Inception-ResNet-V2 architecture, on 85,721 brain MRI scans from 50,876 participants. Applying transfer learning and fine-tuning the model for AD classification, achieved an excellent accuracy of 91.3% in leave-sites-out cross-validation. Here, we examined the performance of DenseNet architecture [12] to classify AD vs healthy controls from two different datasets, and a held-out test dataset. We also used OSA and guided Grad-CAM to improve the interpretability of these models. Guided Grad-CAM provides more detailed visualizations than Grad-CAM, showing specific features rather than just coarse regions. It combines the strength of Grad-CAM – class discrimination – with Guided Backpropagation – fine-grained detail and makes it easier for humans to understand what the model focuses on.

## II. Imaging Data and Preprocessing

The primary dataset for our experiments was the widely-used, publicly available Alzheimer’s Disease Neuroimaging Initiative (ADNI) dataset – a multisite study launched in 2004 to improve clinical trials for the prevention and treatment of AD [13]. We used 4,162 scans for our analysis. The second dataset was from the National Alzheimer’s Coordinating Center (NACC) [14], with a total of 2,347 scans. The third, held-out dataset comes from an Indian population assessed at NIMHANS in Bangalore, India [15,16] – a population not typically well represented in neuroimaging studies. This cohort had 226 participants. The data distribution is shown in **Table 1**; we ensured no overlap between the testing and training data subsets, and the test dataset had only one scan per subject. 3D T1-weighted (T1w) brain MRI volumes were pre-processed using the following steps [17]: nonparametric intensity normalization (N4 bias field correction), ‘skull stripping’ for brain extraction, nonlinear registration to a 3D template – see template details here [18] - with 6 degrees of freedom and isometric voxel resampling to 2 mm. The pre-processed images were of size 91×109×91. The T1w images were scaled using min-max scaling to take values between 0 and 1. For the DenseNet, the image size was further reduced to 80×80×80.

**Table 1.** Data distribution for experiments.

| Dataset | Distribution | Total N | Sex | | Mean Age ± St. Dev. | Diagnosis | |
| --- | --- | --- | --- | --- | --- | --- | --- |
|  |  |  | Male | Female |  | CN | Dementia |
| ADNI | Training Dataset | 3,318 | 1,731 | 1,587 | 75.86 ± 6.74 | 1,950 | 1,368 |
| | Validation Dataset | 433 | 249 | 184 | 75.80 ± 6.94 | 255 | 178 |
| | Test Dataset | 411 | 218 | 193 | 74.15 ± 7.02 | 228 | 183 |
| NACC | Training Dataset | 1,361 | 536 | 825 | 71.15 ± 10.87 | 859 | 502 |
| | Validation Dataset | 321 | 109 | 212 | 70.47 ± 10.57 | 258 | 63 |
| | Test Dataset | 665 | 229 | 436 | 69.68 ± 11.83 | 602 | 63 |
| NIMHANS | Test Dataset | 226 | 125 | 101 | 66.94 ± 7.90 | 128 | 98 |

## III. Deep Learning Architecture, OSA & Grad-CAM

We used the DenseNet architecture [19], as its connectivity pattern promotes efficient feature re-use, incorporating features at multiple scales. The DenseNet architecture is characterized by multiple dense blocks and transition layers. Within a dense block, each layer is connected to all preceding layers, facilitating feature re-use through concatenation: each layer’s output is combined with the feature maps from all prior layers. A typical dense block contains several convolutional blocks, each generating the same number of output channels. Transition layers are employed to manage model complexity and reduce the number of parameters, incorporating convolution and pooling operations to downsample the feature maps. We trained three models -one on ADNI, one on NACC, and one on both datasets combined. Zero-shot classification was also performed on the NIMHANS data. Model performance was assessed using test-balanced accuracy and the F1 score.

Occlusion sensitivity analysis (OSA) is a perturbation-based approach in which input features are altered through occlusion to observe changes in the output. In our case, a cube-shaped mask (7×7×7) is applied, setting the covered voxels to zero. We can monitor the resulting changes in the model’s predictions by systematically moving this mask throughout the entire brain volume (without overlap). The difference in prediction accuracy, measured by the balanced accuracy with and without occlusion, reveals the relative importance of each region. In particular, we examine the change in the classification probability for the class it picked after the final *SoftMax* layer. By mapping these changes, we generate a saliency map highlighting the critical areas the model relies on for its predictions. After generating the OSAs for 100 test subjects, we created a voxel-wise average map to understand any patterns that were consistently used across subjects. In Grad-CAM, the class-discriminative localization map 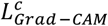 is computed by combining the feature maps *A*^*k*^ from a convolutional layer with weights. These weights 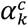 are obtained by global average pooling of the gradients of the target class *c* with respect to the feature map *A*^*k*^. The ReLU activation function is applied to the linear combination of these weighted feature maps to ensure only the positive influences are considered [2].

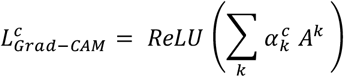

Additionally, *Guided* Grad-CAM includes element-wise multiplication of the Grad-CAM heatmap with Guided Backpropagation to produce high-resolution class-specific visualizations. In the latter, standard backpropagation is performed but with only positive gradients, and the negative gradients are set to zero during backward pass through the ReLu activations. Thus, Guided Grad-CAM combines the spatial information from Grad-CAM with the fine details from Guided Backpropagation, providing a comprehensive visualization tool for understanding deep neural networks decisions. We performed the same steps with Guided Grad-CAM and presented the average group analysis.

## IV. Results

In our experiments, the model trained on a combination of ADNI and NACC data showed better performance on the held-out NIMHANS dataset. The more diverse training data may have provided a more comprehensive representation of the underlying data distribution, enabling the model to generalize better to the holdout dataset; training on a larger combined dataset may also reduce the risk of the model overfitting - learning spurious correlations specific to one dataset. Complementary information present in both datasets may have enabled the model to improve feature representation and selection; combining datasets with differing class distributions may also have balanced the class distribution in the training set, helping the model to distinguish classes. The pooled training set contributed to improved balanced accuracy and F1 Score.

**Figure 1.**
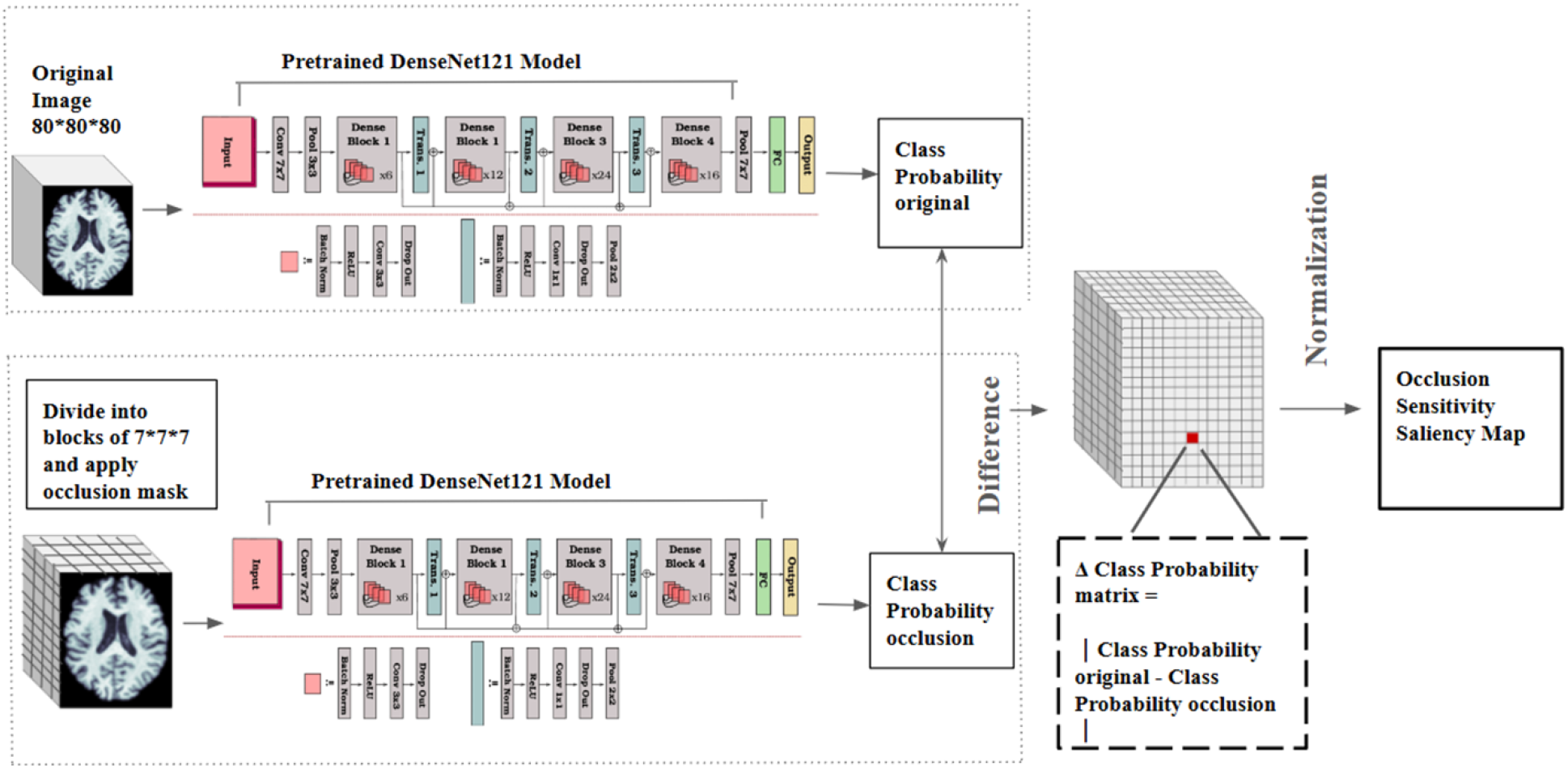
DenseNet Architecture with OSA Model.

**Figure 2.**
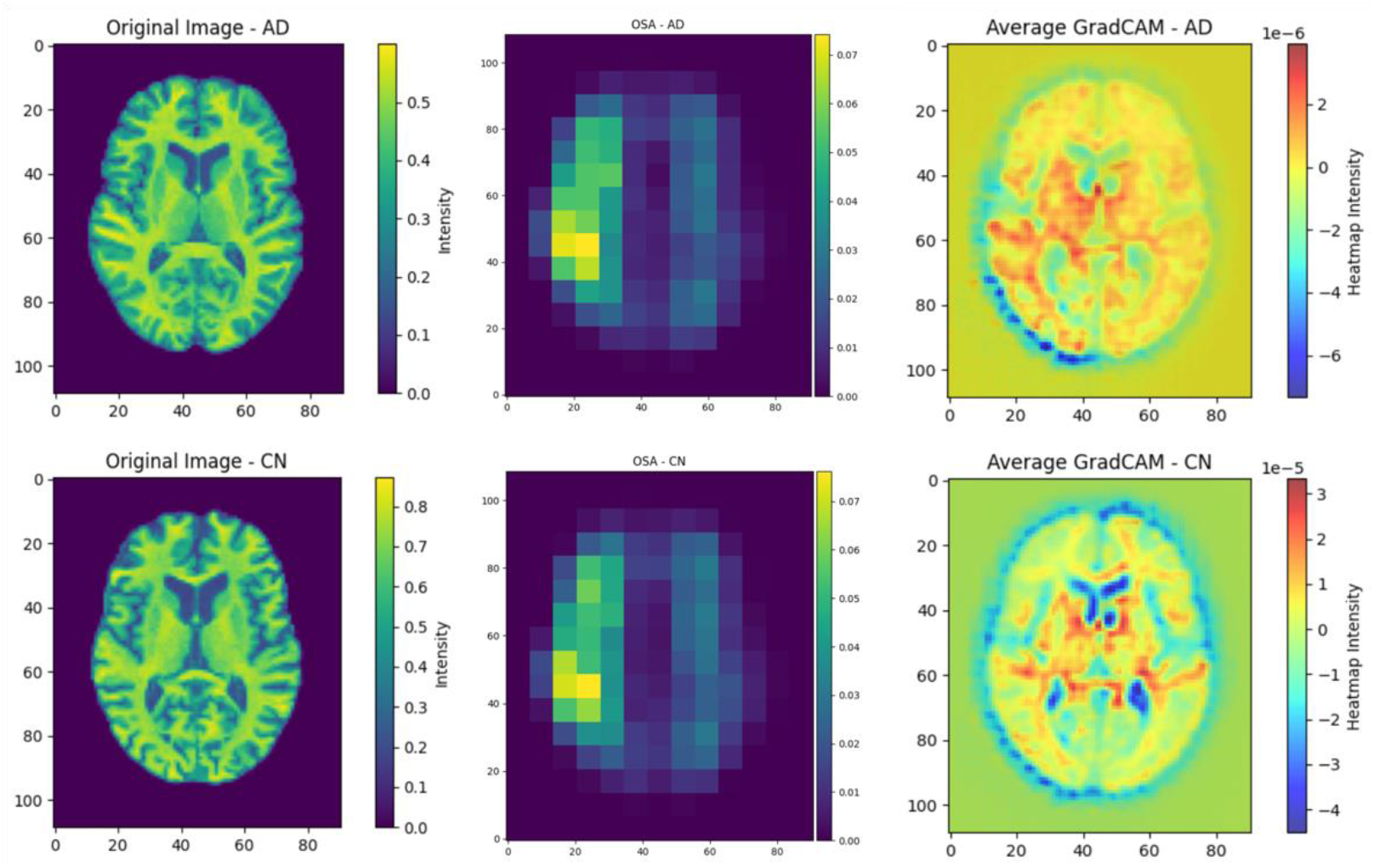
Comparison of explainable models for controls (CN) and participants with dementia in the NACC test dataset. The first column shows the original T1w scan’s middle axial slice (40), for a random subject in the test set. The top row shows a subject with dementia, and the bottom row shows a healthy control participant. The middle column shows the OSA group averages, and the last column shows the corresponding Grad-CAM group averages for 100 participants of the NACC test dataset. Grad-CAM maps detect features at a finer spatial scale than OSA; the OSA maps appear ‘blocky’ as the occlusion kernels do not overlap. If overlap were allowed, these maps would still be spatially smooth. By contrast, Grad-CAM maps show in deep blue the ventricles, which show strong group differences between patients and controls.

The OSA analysis for the NACC dataset was performed by using all subjects from the test dataset and visualizing features used by the model. The delta probability matrix was computed for each subject and the group average was calculated to visualize disease patterns. As the absolute value of the probability delta was calculated, the change is reflected as a positive delta in each case. Grad-CAM was similarly computed for each subject in the test dataset, and the group average calculated, for comparison with OSA. The Grad-CAM maps show great spatial detail and better feature localization than OSA. A lower mask size for OSA may improve the spatial detail in the OSA maps. Comparing it to **Figure 3**, the pattern is congruent with the AD effect found using other methods to map the atrophy profile, such as TBM [20].

**Table 2.** Classification results for the three models. B.A. indicates balanced accuracy and F1 indicates F1 Score. The NIMHANS dataset is treated as a zero-shot test dataset. The left rows are the training datasets, and the top columns are the test datasets respectively (if the same dataset appears as test data, a subset was used without subject overlap). Bold indicates the best balance accuracy on NIMHANS test set.

|  | Training Datasets | Testing Datasets |  |  |  |  |  |
| --- | --- | --- | --- | --- | --- | --- | --- |
|  |  | NACC |  | ADNI |  | NIMHANS |  |
|  |  | B.A. | F1 | B.A. | F1 | B.A. | F1 |
|  | NACC | 0.79 | 0.81 | 0.76 | 0.76 | 0.78 | 0.75 |
|  | ADNI | 0.76 | 0.65 | 0.78 | 0.76 | 0.80 | 0.76 |
|  | NACC + ADNI | 0.76 | 0.76 | 0.78 | 0.73 | **0.87** | 0.85 |

**Figure 3.**
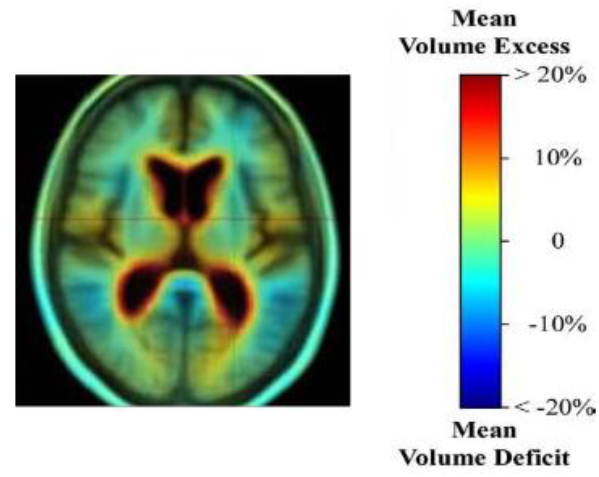
Atrophy map comparing an AD patient group to healthy controls, using tensor-based morphometry in the ADNI dataset. Although maps of salient features are hard to validate without ground truth on the location of the disease, complementary image analysis methods, such as TBM, show ventricular expansion and subcortical atrophy. These key signs of dementia are used by radiologists for diagnosis. (Reproduced from Hua et al. [20]).

## V. Conclusions and Future Work

Our experiments examined how well models trained on datasets of mostly North American and European ancestry generalize to a new dataset from Indian participants. Our findings suggest that adding more data while training may improve performance on holdout test sets. We also used occlusion sensitivity analysis and Grad-CAM to visualize features used by the deep learning models that were important for classification. Future work will evaluate more complex explainable AI models such as attention flow and attention rollout, which are typically used for vision transformers. We will also evaluate more advanced imaging modalities including diffusion-weighted MRI [21], to identify important features for multimodal diagnostic classification.

## Acknowledgments

This work was supported by NIH grant R01 AG060610 funded by the National Institute on Aging (NIA) and the Fogarty International Center (FIC), as well as by NIH NIA grant U01 AG068057 (‘AI4AD’). NIMHANS data collection was supported by the Department of Science and Technology, Govt. of India, grant nos. DST-SR/CSI/73/2011 (G), DST-SR/CSI/70/2011 (G) and DST/CSRI/2018/249 (G).

## Notes

### Competing Interest Statement

The authors have declared no competing interest.

## References

[1] New AI legislation’s reach extends into European Healthcare. Osborne Clarke. (2024). https://www.osborneclarke.com/insights/new-ai-legislations-reach-extends-european-healthcare.

[2] Selvaraju, R. R., et al. (2017). Grad-CAM: Visual explanations from deep networks via gradient-based localization. In Proc. IEEE International Conference on Computer Vision (pp. 618–626).

[3] Zhang, Y., et al. (2021). Grad-CAM helps interpret the deep learning models trained to classify multiple sclerosis types using clinical brain magnetic resonance imaging. NeuroImage: Clinical, 30, 102642.

[4] Qin, C., et al. (2021). A large-scale multimodal neuroimaging dataset to identify brain disorders based on anatomical and functional markers. Scientific Data, 8(1), 1–21.

[5] Sarraf, S., & Tofighi, G. (2022). Classification of Alzheimer’s disease using 3D convolutional neural networks and explainable artificial intelligence. Informatics in Medicine Unlocked, 26, 100732.

[6] Simonyan, K., Vedaldi, A., & Zisserman, A. (2013). Deep inside convolutional networks: Visualising image classification models and saliency maps. arXiv preprint arXiv:1312.6034.

[7] Zeiler, M. D., & Fergus, R. (2014). Visualizing and Understanding Convolutional Networks. European Conference on Computer Vision (ECCV).

[8] Moguilner, S., et al. (2023). Visual deep learning of unprocessed neuroimaging characterises dementia subtypes and generalises across non-stereotypic samples. EBioMedicine, 90.

[9] World Health Organization, “Dementia,” 2022. https://www.who.int/news-room/fact-sheets/detail/dementia.

[10] Schlemper, J., et al. (2019). Attention gated networks: Learning to leverage salient regions in medical images. Medical Image Analysis, 53, 197–207.

[11] Lu, B., Li, H. X., et al. (2022). A practical Alzheimer’s disease classifier via brain imaging-based deep learning on 85,721 samples. Journal of Big Data, 9(1), Article 101.

[12] T. Zhou et al., (2022). Dense convolutional network and its application in medical image analysis. BioMed Research International, 2022, 1–22.

[13] Veitch D., et al., “Understanding disease progression and improving Alzheimer’s disease clinical trials: Recent highlights from the Alzheimer’s Disease Neuroimaging Initiative,” Alzheimer’s Dement. 15(1):106–152 (2019).

[14] National Institute on Aging. (2005). National Alzheimer’s Coordinating Center (NACC) Database [Data set]. https://www.alz.washington.edu.

[15] Bharath, S., et al. (2017). A Multimodal Structural and Functional Neuroimaging Study of Amnestic Mild Cognitive Impairment. American Journal of Geriatric Psychiatry, 25(2), 158–169.

[16] Joshi, H., et al. (2019). Differentiation of Early Alzheimer’s Disease, Mild Cognitive Impairment, and Cognitively Healthy Elderly Samples Using Multimodal Neuroimaging Indices. Brain Connectivity, 9(9), 730–741.

[17] Lam, P., et al., “3-D Grid-Attention Networks for Interpretable Age and Alzheimer’s Disease Prediction from Structural MRI,” arXiv (2020).

[18] Zhu, A.H., et al. (2021). ‘Age-Related Heterochronicity of Brain Morphometry may Bias Voxelwise Findings.’ IEEE 18th International Symposium on Biomedical Imaging (ISBI), pp. 836–839.

[19] Huang, G., et al., “Densely Connected Convolutional Networks,” CVPR, 4700–4708 (2017).

[20] Hua, X., et al. (2008). Tensor-based morphometry as a neuroimaging biomarker for Alzheimer’s disease: An MRI study of 676 AD, MCI, and normal subjects. NeuroImage, 43, 458–469.

[21] Chattopadhyay, T., et al., (2023). “Predicting dementia severity by merging anatomical and diffusion MRI with deep 3D convolutional neural networks.” In the 18th International Symposium on Medical Information Processing and Analysis (Vol. 12567, pp. 90–99). SPIE.

